# Urethral luminal epithelia are castration-insensitive progenitors of the proximal prostate

**DOI:** 10.1101/2020.02.19.937615

**Authors:** Diya B. Joseph, Gervaise H. Henry, Alicia Malewska, Nida S. Iqbal, Hannah M. Ruetten, Anne E. Turco, Lisa L. Abler, Simran K. Sandhu, Mark T. Cadena, Venkat S. Malladi, Jeffrey C. Reese, Ryan J. Mauck, Jeffrey C. Gahan, Ryan C. Hutchinson, Claus G. Roehrborn, Linda A. Baker, Chad M. Vezina, Douglas W. Strand

## Abstract

Castration-insensitive epithelial progenitors capable of regenerating the prostate are concentrated in the proximal region close to the urethra, but the identification of these cells has been limited to individual cell surface markers. Here, we use single cell RNA sequencing (scRNA-seq) to obtain a cellular anatomy of the mouse prostate and urethra and create a comparative map with the human. These data reveal that previously identified facultative progenitors marked by TROP2, Sca-1, KRT4, and PSCA are actually luminal epithelia of the urethra that extend into the proximal prostate. These mouse urethral cells are the human equivalent of previously identified club and hillock urethral cells. Castration decreases androgen-dependent prostate luminal epithelia as expected, but TROP2+ urethral luminal epithelia survive and expand into the prostate. Benign prostatic hyperplasia (BPH) has long been considered an ‘embryonic reawakening’, but the cellular origin of peri-urethral growth is unclear. We use scRNA-seq and flow cytometry to demonstrate an increase in urethral luminal epithelia within glandular nodules from patients with BPH, which are further enriched in patients treated with a 5 alpha reductase inhibitor. These data demonstrate that the putative prostate progenitors enriched by castration in the proximal prostate are an expansion of urethral luminal epithelia and that these cells may play an important role in the etiology of human BPH.

**Significance Statement:** The prostate involutes after castration, but regrows to its original size with androgen replenishment. This observation prompted the search for a castration-insensitive prostate progenitor. Here, Joseph *et al*. produce a comparative cellular atlas of the prostate and urethra in the mouse *vs.* human, discovering an equivalent urethral luminal epithelial cell type that extends into the proximal prostatic ducts and expresses previously identified markers of facultative prostate progenitors. Urethral luminal epithelia are established before prostate budding in human and mouse development, and expand after castration in the mouse and after 5 alpha reductase inhibitor treatment in human BPH. These data suggest that luminal epithelia of the urethra are castration-insensitive cells of proximal ducts that may act as progenitors in human BPH.

## Introduction

The mouse and human prostate develop as a series of solid epithelial buds from the primitive urethra, or urogenital sinus. Buds undergo a period of branching morphogenesis and canalization resulting in a tree-like network of proximal ducts and distal glands (1-3). We previously produced a cellular atlas of the young adult human prostate and prostatic urethra using single cell RNA sequencing (scRNA-seq) on proximal and distal anatomical regions. These data confirmed known cell types of the prostate such as basal, luminal, and neuroendocrine epithelia and led to the identification of previously unknown hillock and club epithelia as the major luminal epithelial cell types composing the prostatic urethra and its proximal ducts (4). ‘Intermediate’ cells labeled with Keratin 19, Keratin 8/18, and Keratin 5/14 are said to exist transiently in the adult and prior to prostate budding from the urogenital sinus during development (5, 6). Prominent studies in other organs have shown that a rigorous mapping of equivalent mouse cell types with objectively defined cell type-specific markers can lead to better models of human disease (7, 8), but a similar comparative cellular atlas of the human and mouse prostate has not been compiled.

The human prostate consists of a network of 12-14 paired proximal ducts located in the transition zone that empties secretory products of distal glandular units into the urethra (9). Benign Prostatic Hyperplasia (BPH) is an expansion of the transition zone surrounding the urethra that occurs in a majority of aging men and is commonly treated with 5 alpha reductase inhibitors (5ARI) to cause apoptosis of androgen-dependent luminal epithelia (10, 11). The cellular origin of peri-urethral prostate growth is unknown, but has been hypothesized to be an embryonic reawakening (12-15).

During development, multipotent basal epithelial progenitors give rise to basal, luminal, and neuroendocrine cell types (6, 16). In the adult, basal and luminal lineages are unipotent (17), but an ‘intermediate’ cell type with basal and luminal characteristics is said to give rise to luminal cells as well (6). The prostate displays potent regenerative capacity after castration and subsequent androgen replenishment (18), prompting the search for castration-insensitive prostate progenitors. Lineage tracing of luminal epithelia with Keratin 8 demonstrates that, after castration and regeneration, prostate luminal cells are largely derived from unipotent luminal progenitors (17, 19, 20). Several studies have identified cell populations characterized by expression of BMI1 (21), Sca-1 (22, 23), TROP2 (24), LY6D (25), and KRT4 (26) that are enriched in the proximal region of the mouse prostate after castration, form spheroids at high frequency, and can act as facultative progenitors in prostate regeneration assays. A proximal to distal expansion of progenitors in human lineage tracing studies has also been demonstrated (27, 28). Proximal epithelia lineage traced with either basal (Keratin 5/14) or luminal (Keratin 8/18) markers or purified with individual cell surface markers show progenitor activity (22, 29-31), but these transgenes and cell surface markers are expressed across multiple tissue types, making it difficult to precisely capture the identity of cell types of interest. These issues exemplify the need for objective transcriptional identities and cell surface markers of each cell type in the lower urinary tract.

Cell identity has historically been defined by individual markers, and the interpretation of data from experimental mouse models relies on these definitions. Here we perform scRNA-seq on the mouse prostate and urethra to build a comparative cellular atlas to the human in order to bolster the molecular and anatomical definition of each cell type (4, 32). We use scRNA-seq, flow cytometry, and immunohistochemistry to demonstrate that the proximal castration-insensitive progenitor is a urethral luminal epithelial cell type that exists in the urogenital sinus before prostate budding and extends into the ductal network of the proximal prostate in the normal adult. Urethral luminal epithelia are increased in the proximal prostate of castrated mice as well as in human BPH patients treated with 5ARIs. When comparing the transcriptome of urethral luminal epithelia in normal young adults vs. BPH patients, a pronounced immunomodulatory phenotype is observed. In summary, our data demonstrate that luminal progenitors in the proximal prostate are actually an extension of castration-insensitive, immunomodulatory urethral luminal epithelia and that this cell type is resistant to 5ARI treatment in human BPH.

## Results

### Identification of equivalent epithelial cell types of the prostate and urethra in the mouse vs. human

We set out to identify equivalent human and mouse epithelial cell types in the prostate and urethra by scRNA-seq, given the essential role of the mouse in modeling human prostate disease. First, we combined ventral, dorsolateral, and anterior mouse prostate lobes for digestion and barcoding in aggregate. We also independently barcoded cells from the dissected urethra, which contains ejaculatory ducts and the epithelium of the prostatic urethra enclosed by the skeletal muscle of the rhabdosphincter. The data from the prostate and urethral regions were aggregated (Figure S1A) and equivalent cell types were identified by bioinformatical comparison to our previously published human prostate and urethra scRNA-seq dataset (4). After labeling major cell lineages (leukocytes, endothelia, fibromuscular stroma, and epithelia) in the mouse data using our previous human data (Figure S1A, B), we sub-clustered the epithelia and displayed clusters split by anatomical region (Figure 1A, B).

**Figure 1.**
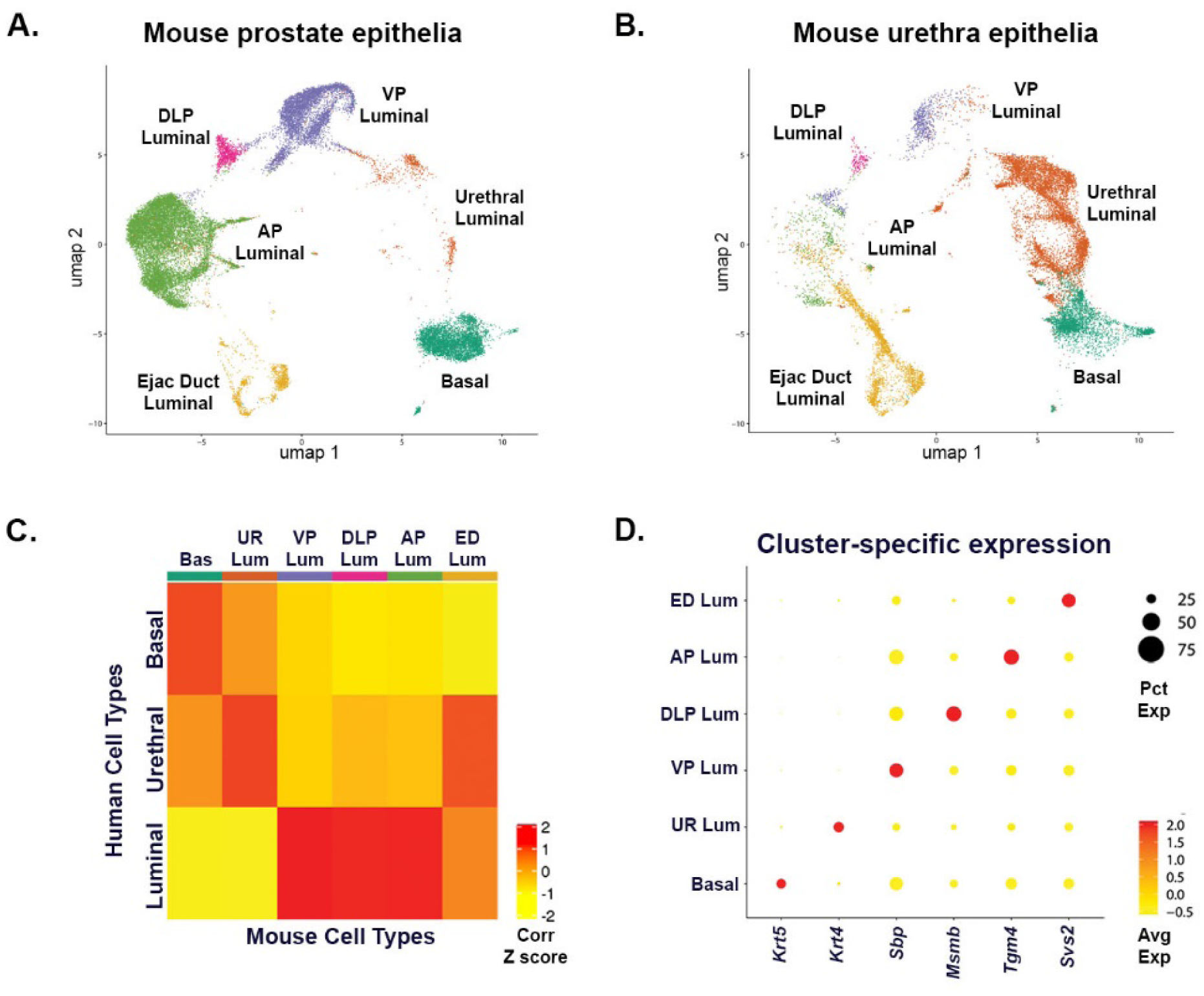
Identification of epithelial cell types of the mouse prostate and prostatic urethra. **(A, B)** Mouse prostate lobes (n=4 mice) were dissected away from the rhabdosphincter of the urethra (n=3 mice) and each anatomical region was processed into a single cell suspension and barcoded separately for scRNA-seq. The data were aggregated and sub-clustered by epithelial lineage (see Figure S1), and separated into prostate **(A)** and urethra **(B). (C)** Statistical correlation of each epithelial cluster of the mouse prostate and urethra with human epithelial cell types. **(D)** Dot plot of differentially expressed genes for each cluster. AP, anterior prostate; VP, ventral prostate; DLP, dorsolateral prostate; ED, Ejaculatory duct; UR, urethra; Bas, basal epithelia; Lum, luminal.

Mouse epithelial cell types were defined based on correlation to basal, luminal, and urethral epithelia in the human (4) and included one basal, two urethral, and three luminal clusters (Figure 1C, Figure S1C, D). Given the enrichment of *Pax2* in Wolffian duct derivatives (33), we labeled one of the urethral clusters as ejaculatory duct (ED). The three clusters of luminal epithelia corresponded to specific prostate lobes based on previous studies that identified lobe-specific expression of *Msmb* (dorsolateral, DLP), *Tgm4* (anterior, AP), and *Sbp* (ventral, VP) (34). Figure 1D demonstrates the enrichment of differentially expressed genes (DEGs) for each basal and luminal epithelial cluster. The full list of significant DEGs expressed in discrete cell types of the mouse prostate and urethra can be found in Dataset S1. Sequencing metrics for scRNA-seq on mouse prostate and urethra can be found in Dataset S2.

A novel mouse urethral luminal epithelial cell type was identified based on correlation to human club and hillock urethral cell types. Mouse urethral cells highly expressed Keratin 4 (Figure 1D) and were enriched in the urethra single cell preparation compared to the prostate (Figure 1A, B). We were not able to identify a stereotypical club urethral epithelial cell type (characterized by expression of secretoglobins), in the mouse either in the scRNA-seq data or by immunostaining (Figure S2). However, mouse urethral luminal cells share properties of both human club and hillock epithelia at the pan-transcriptomic level including expression of *Psca* (Figure S2B) (4).

### Urethral luminal epithelia extend into the proximal ducts of the prostate and express markers of prostate progenitor and intermediate cells

Proximal ducts of the mouse prostate have been shown to contain a concentration of androgen-independent progenitors (22, 29-31), but identification and purification of these cells has relied on individual markers. To determine precisely where luminal epithelia of the urethra are located, we performed immunofluorescent staining on transverse whole mount sections with antibodies to KRT4 (urethral luminal), KRT5 (basal epithelia of urethra and prostate), and NKX3.1 (prostate luminal) (35). We observed that luminal epithelia of the urethra and proximal ducts within and leading out of the rhabdosphincter are KRT4+/NKX3.1- and transition immediately into KRT4- /NKX3.1+ prostate epithelial cells (Figure 2A). The transition from urethral identity to prostate identity in human proximal ducts can be similarly highlighted using KRT13 (urethral luminal cell) and ACPP (prostate luminal cell) (Figure 2B). Similar to our previous human prostate study, neuroendocrine epithelia are an extremely rare cell type that do not cluster independently, but can be identified using known markers such as CGRP and CHGA (Figure S3A, B). We also observed that neuroendocrine epithelia are enriched within the urethra and proximal prostatic ducts in the mouse and human (Figure S3C, D).

**Figure 2.**
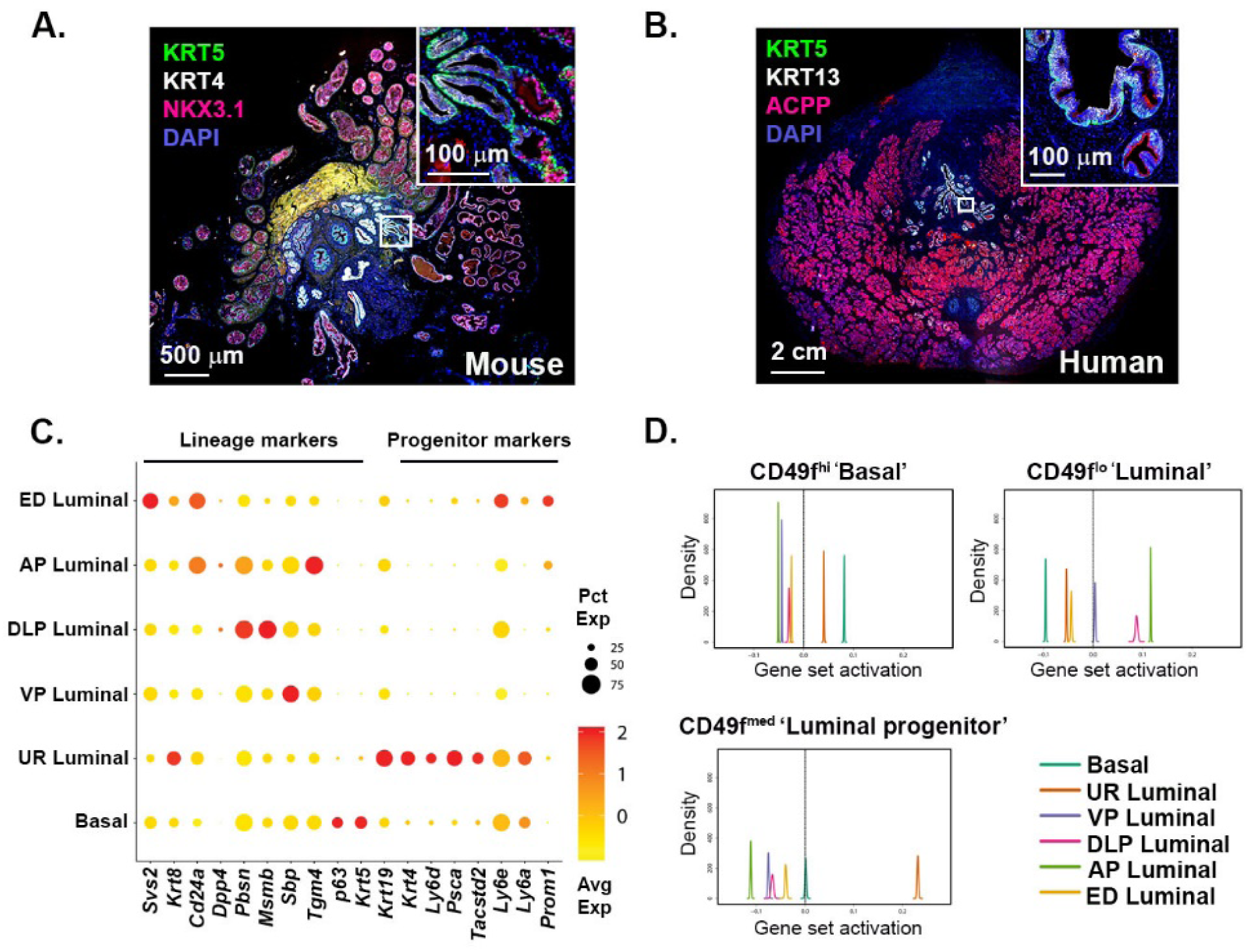
Urethral luminal epithelia display putative progenitor markers and extend into the proximal prostate after castration. **(A)** KRT4 labeled mouse urethral luminal epithelia within the urethra and proximal prostate ducts transitions into NKX3.1+ prostate secretory luminal epithelia ducts (n=3 mice). **(B)** KRT13 labeled human urethral luminal epithelia within the prostatic urethra transitions into ACPP+ prostate secretory luminal epithelia in the transition zone (n=3 human samples). **(C)** Dot plot of mouse prostate and urethral epithelial cell type-specific markers and putative stem cell markers. **(D)** Correlation of mouse scRNA-seq clusters to transcriptomic signatures of basal, luminal, and LSC^med^ ‘luminal progenitors’ from (26). AP, anterior prostate; VP, ventral prostate; DLP, dorsolateral prostate; ED, Ejaculatory duct; UR, urethra.

According to previous studies, epithelia that are enriched in facultative stem cell assays display markers such as BMI1 (21), Sca-1 (22, 23), TROP2 (24), LY6D (25), and KRT4 (26). A dot plot of putative stem cell markers demonstrates that these markers are almost exclusively expressed by the urethral luminal epithelial cluster, although some proposed stem cell markers such as CD133 (*Prom1*) are enriched in luminal epithelia of the ejaculatory ducts (Figure 2C). Krt19 has historically been associated with KRT5+/KRT8+ ‘intermediate’ cells transitioning from basal to luminal (5), but is highly enriched in urethral luminal epithelia. CD49f and Sca-1 are commonly used to isolate prostate basal (CD49f^hi^/Sca-1^hi^) and luminal (CD49f^lo^/Sca-1^lo^) epithelia (24, 26, 31). Multiple studies have also identified a progenitor ‘LSC^med^’ population (CD49f^med^/Sca-1^hi^) enriched in castration and tumorigenesis (22, 26). Correlation analysis of transcriptomic signatures from cells in each of these three flow cytometry gates from Sackmann Sala *et al*. (26) versus each of our scRNA-seq epithelial clusters demonstrates that the urethral luminal epithelia cluster is highly similar to the LSC^med^ androgen-independent luminal progenitor (Figure 2D). These data suggest that the full identity of the luminal prostate progenitor that is increased in castration and prostate tumors is actually the urethral luminal cell.

### TROP2+ urethral luminal epithelia are enriched in castration

Given the transcriptional similarity of urethral luminal epithelia to facultative progenitors (Figure 2D), we set out to quantitate the frequency of urethral luminal epithelia in discrete anatomical regions of an intact mouse. We dissected the urethra (UR) away from proximal and distal prostate regions for digestion into single cells (Figure 3A) and developed a flow cytometry scheme using TROP2 (*Tacstd2*) as a cell surface marker, given its selectivity for urethral luminal epithelia over Sca-1 (*Ly6a/e*) in the scRNA-seq data (Figure 2C). However, TROP2 and Sca-1 label a largely overlapping population of cells (data not shown).

**Figure 3.**
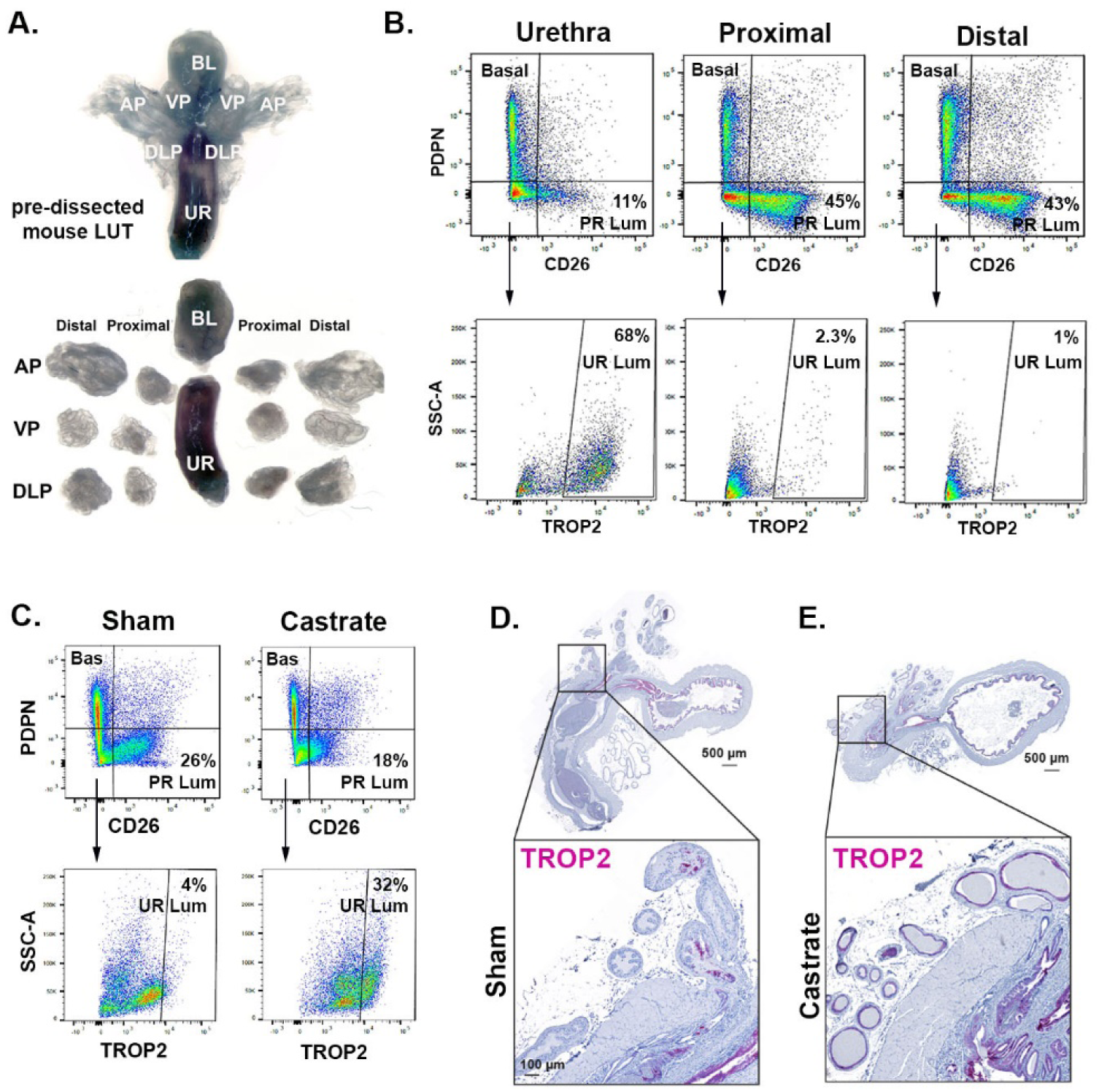
TROP2+ urethral luminal epithelia are enriched in the prostates of castrated mice. **(A)** Dissection of mouse prostate lobes away from urethra, and into proximal and distal regions. **(B)** Flow cytometry of CD26+ prostate luminal epithelia, PDPN+ basal epithelia, and TROP2+ urethral epithelia in the urethra, proximal prostate, and distal prostate. Data representative of n=4 independent experiments. **(C)** Quantitation of flow cytometry results show that TROP2+ urethral luminal epithelia are enriched in prostates of castrated mice. **(D, E)** Luminal epithelia in the prostatic urethra, the bladder, and proximal prostatic ducts are marked by TROP2 and expands in castrated mice. Images representative of n=3 mice per group. LUT, lower urinary tract; AP, anterior prostate; VP, ventral prostate; DLP, dorsolateral prostate; BL, bladder; UR, urethra; Bas, basal epithelia; PR Lum, prostate luminal epithelia; UR Lum, urethral luminal epithelia.

Our gating scheme first separates leukocytes (CD45+), epithelia (CD326+), and stroma (CD45-/CD326-). CD326+ epithelia are then gated into prostate luminal (CD26+) and non-prostate luminal (CD26-) cells. Previously used cell surface markers such as CD24 (*Cd24a*) label prostate luminal epithelia (22), but also luminal epithelia of the ejaculatory ducts and urethra, whereas CD26 (*Dpp4*) is selectively enriched in prostate luminal epithelia (Figure 2C). CD26+ prostate luminal epithelia are enriched in the proximal and distal prostate (42.1 ± 4.8% and 40.1 ± 4.4% of total epithelia, respectively) compared to the urethra (12.5 ± 2.1% of total epithelia). Because basal epithelia can also be positive for TROP2, urethral luminal epithelia are more accurately identified in the PDPN-/CD26- (non-basal, non-prostate luminal) gate. TROP2+ urethral luminal cells represent 20.6 ± 1.6% of total epithelia within the urethral region, and 0.44 ± 0.10% of total epithelia within the proximal prostate, compared to 0.22 ± 0.05% in the distal prostate (from 4 independent experiments) (Figure 3B). Figure S4 demonstrates that KRT4 and TROP2 identify the same cell type by displaying overlapping expression in the mouse urethra and proximal prostatic ducts.

Proximal KRT4+/Sca-1+/NKX3.1-prostate progenitors have been shown to be enriched after castration as well as in tumorigenesis (23, 26, 31). To determine whether TROP2+ urethral luminal epithelia are likewise enriched after castration, we used our optimized flow cytometry antibody panel to quantitate epithelial cell types in castrated *vs.* sham castrated mice. TROP2+ urethral luminal epithelia are increased in prostates from castrated mice (11.9% of total epithelia) compared to sham castrated mice (1.33% of total epithelia) animals, while CD26+ prostate luminal epithelia represent 26% of total epithelia in sham mice compared to 18% in castrated mice (Figure 3C). These data are highly similar to the 4-fold increase in Sca-1+/CD49f^med^ luminal epithelia (‘LSC^med^’) after castration in a previous study and the transcriptomic signature of this population was highly correlated with our urethral luminal signature (Figure 2D), further suggesting these are the same cell type (26). We demonstrate *in situ* that the increased percentage of TROP2+ urethral luminal epithelia after castration is due to survival and expansion into the prostate using a TROP2 antibody in sham and castrate whole mount lower urinary tracts (Figure 3D, E).

### Urethral identity is established prior to prostate budding in mouse and human

The prostate goes through a developmental process of bud initiation, branching morphogenesis, and canalization (1, 2). Although several adult epithelial cell populations can act as facultative progenitors in tissue regeneration and spheroid assays (17, 36), lineage tracing analyses suggest that the basal and luminal lineages are largely unipotent in the adult (6, 17). However, the transgenes used in these studies (*Krt5* and *Krt8*) label basal and luminal epithelial cells from prostate, urethra, and ejaculatory ducts, highlighting the need for developing tissue-specific markers and understanding when and where they are expressed during development.

To determine whether the identity of urethral epithelia is established early in the developmental process, we examined developmental stages of the mouse and human prostate. Mouse urethral luminal epithelia could be identified by KRT4 during the earliest stages of bud initiation (E18.5) where these cells occupied the inner region of prostate buds. During the branching stage (P9), KRT4+ urethral luminal cells were present in the urethra and branching prostate ducts (Figure 4A, B) (1, 2). In the adult mouse prostate, KRT4+ urethral cells were restricted to the urethra and urethra-proximal ductal segments (Figure 4C). In the human male, prostate buds initiate at 10-13 weeks and undergo branching morphogenesis beginning at 12-18 weeks (37). At 12 weeks gestation, club and hillock cells marked by SCGB1A1 and KRT13, respectively, could be observed in the urethra and developing buds (Figure 4D). At 19 weeks gestation, club and hillock luminal epithelia could be observed in the urethra and in proximal segments of branching prostatic ducts (Figure 4E). In the adult human male, club and hillock urethral luminal epithelia extend into the proximal ductal network as shown in the sagittal plane (Figure 4F). These results demonstrate that the identity of urethral cell types is established before prostate budding and branching morphogenesis, which could have implications for human BPH if it is indeed a reawakening of embryonic processes. Replicates for each stain on independent specimens can be found at https://doi.org/10.25548/16-WM8C.

**Figure 4.**
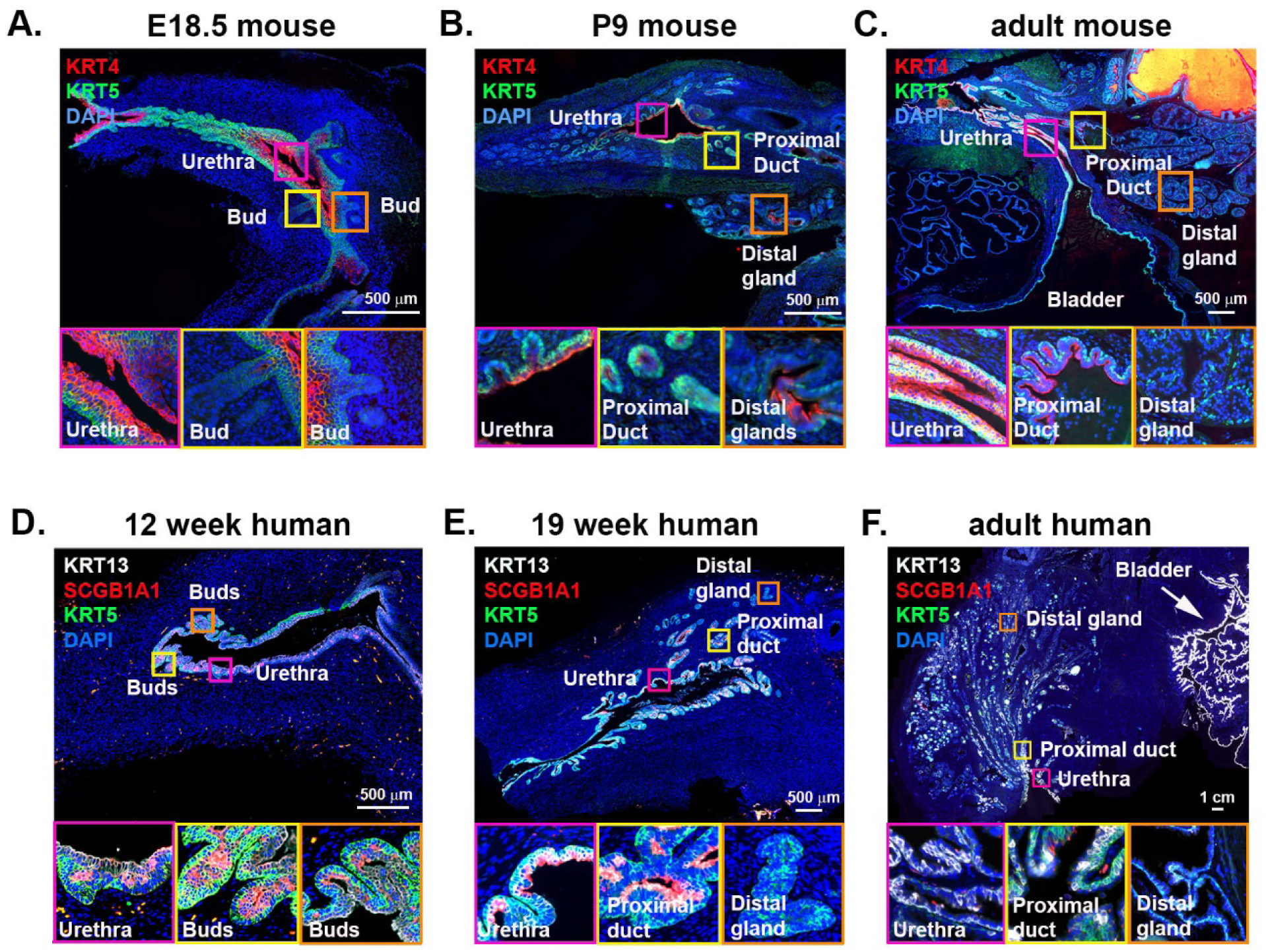
Urethral epithelial identity is established early in prostate budding. Mouse lower urinary tract sections were labeled with antibodies to KRT4 (urethral luminal epithelia) and KRT5 (basal epithelia). Stages shown are **(A)** Embryonic day 18.5 (budding) **(B)** Postnatal day 9 (branching) and **(C)** Adult mouse prostate. Human lower urinary tract sections were labeled with antibodies to KRT13 (hillock epithelia), SCGB1A1 (club epithelia) and KRT5 (basal epithelia). Stages shown are **(D)** 12 weeks gestation (budding) **(E)** 19 weeks gestation (branching) and **(F)** adult human prostate.

### Urethral epithelial cells are increased in human BPH and are resistant to 5ARI treatment

Benign prostatic hyperplasia (BPH) is a continuous expansion of the prostate transition zone that affects 70% of men over 70 years old and has been hypothesized to be a reawakening of developmental processes (14, 15). To identify cell types present in BPH, we performed scRNA-seq on glandular nodules from three patients who underwent simple prostatectomy for lower urinary tract symptoms (see Dataset S2 for sequencing metrics and Dataset S3 for clinical data on each patient) and sub-clustered the epithelia for comparison to three young normal prostates (4) (Figure 5A). The results demonstrate an increase in club urethral luminal epithelia from 2% of epithelia in the normal prostate to 9% of epithelia within glandular BPH. We previously established PSCA as a highly specific cell surface marker capable of capturing a 94% pure population of prostatic urethral luminal epithelia in the human (4). To confirm whether PSCA+ urethral luminal epithelia are increased in glandular BPH in a larger cohort, we performed flow cytometry on non-diseased prostates from young organ donors (n=6) for comparison to glandular BPH specimens from men (n=6) undergoing simple prostatectomy (Dataset S3, Figure S5). PSCA+ urethral luminal epithelia are significantly increased in glandular BPH compared to young donor prostates (Figure 5B). With the aggregation of normal and BPH scRNA-seq data, we generated cell type-specific DEGs to get a basic understanding of the functional contributions of each cell type in disease. Dataset S5 displays the significant DEGs for each epithelial cell type in normal vs. BPH. To annotate the functional changes in club cells in BPH *vs*. normal human prostate, we performed a KEGG analysis (38) of DEGs in BPH clubs cells compared to normal club cells. In the normal adult human, urethral luminal epithelia are highly enriched in metabolic pathways (4), which is the same for the mouse (data not shown). However, Figure 5C shows that the majority of the genes that are significantly upregulated in club cells from BPH tissue belong to immunomodulatory pathways.

**Figure 5.**
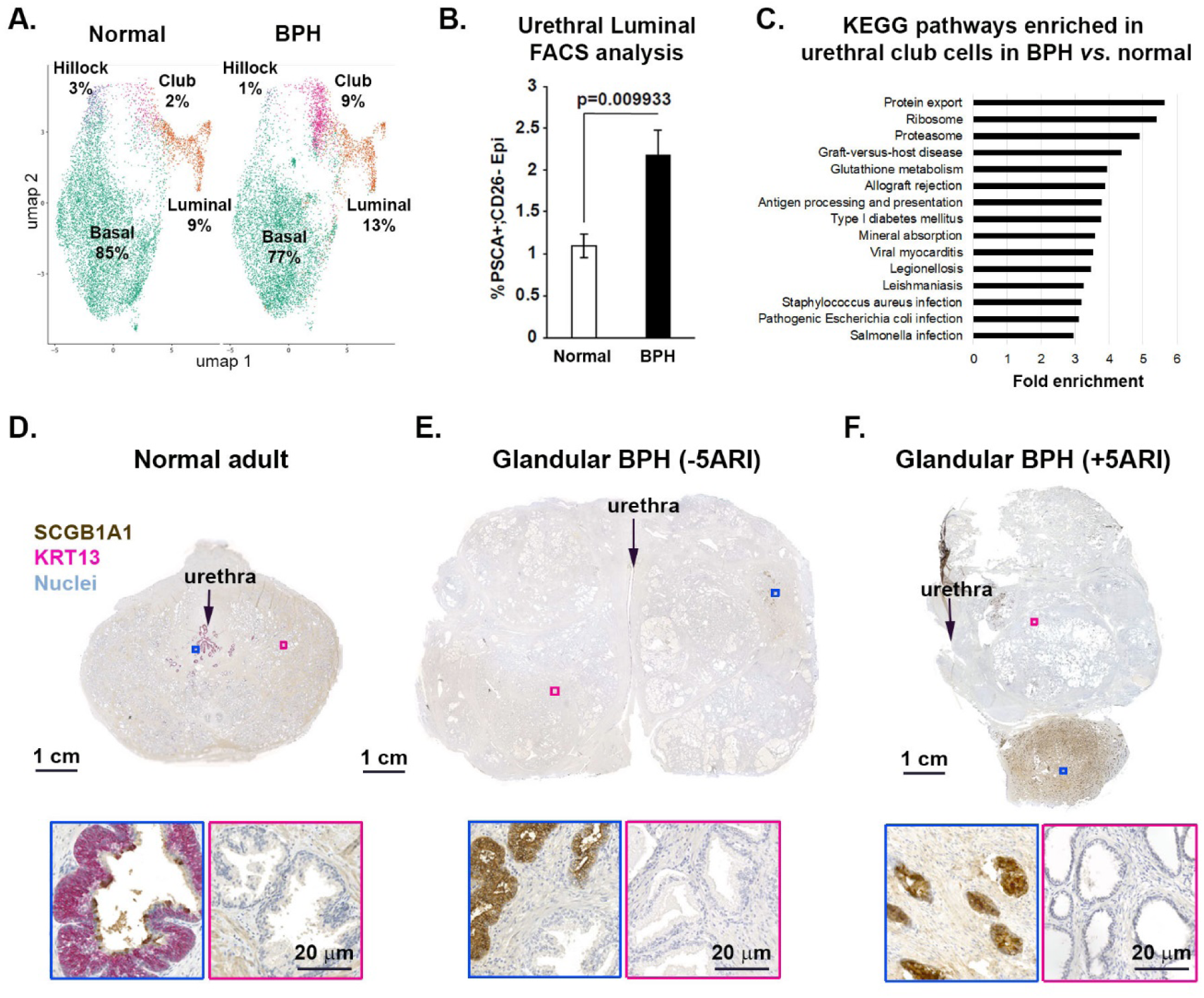
Urethral epithelia are enriched in human BPH and are resistant to 5ARI treatment. **(A)** scRNA-seq of tissue from 3 patients with glandular BPH demonstrates an enrichment of club epithelia compared to the 3 young adult normal prostates. **(B)** PSCA+ urethral epithelia are enriched in glandular BPH compared to normal prostate using flow cytometry (n=6 per group). **(C)** Fold enrichment of top 15 KEGG pathways significantly upregulated in club urethral luminal cells from BPH vs. normal prostate. **(D-F)** Dual IHC for KRT13 and SCGB1A1 demonstrates an expansion of urethral epithelia in glandular BPH (n=10) and a further enrichment in 5ARI-treated patients (n=11) compared to young normal adult prostate (n=5).

Five alpha reductase inhibitors (5ARIs) are widely used to treat BPH patients. 5ARI treatment shrinks prostate volume through luminal cell apoptosis by inhibiting the conversion of testosterone to dihydrotestosterone (DHT) (10). Given that castration in the mouse enriched for TROP2+/KRT4+ urethral luminal epithelia (Figure 3C-E), we set out to test whether urethral epithelial cells were increased in areas of atrophy within human BPH in patients treated with a 5ARI. Accordingly, we performed dual immunohistochemistry for club (SCGB1A1) and hillock (KRT13) urethral luminal epithelia in young men and in men with BPH that were not taking a 5ARI. We could detect rare SCGB1A1+ club cells in 5ARI-naïve men with BPH similar to the scRNA-seq data (Figure 5D, E). However, analysis of whole mounted prostates from 11 different 5ARI-treated men revealed a striking increase in the number of club and hillock epithelia in focal areas of atrophy, further suggesting DHT independence of urethral luminal epithelia in areas of prostate luminal cell apoptosis (Figure 5F, Dataset S3). Replicates for each stain on independent specimens can be found at https://doi.org/10.25548/16-WM8C.

## Discussion

We have produced a cellular anatomy of the mouse prostate and urethra, by comparing the molecular identity of equivalent cell types and their locations to the human (Figures 1 and 2). We demonstrate that many of the markers previously used to characterize castration-insensitive prostate progenitors (20, 22-26, 29-31, 39-43) are highly expressed by urethral luminal epithelia, suggesting that the facultative progenitors of the proximal prostate described as NKX3.1-luminal epithelia are of urethral origin (Figure 2C). Moreover, similar to the human anatomy, mouse urethral luminal epithelia extend into the proximal ducts of the prostate before transitioning to prostate epithelial identity (Figure 2A, B), which explains why NKX3.1-/Sca-1+/TROP2+ cells are enriched in the proximal prostate (22, 24, 31).

The characterization of prostate epithelial cells that survive castration and serve as cells of origin for regeneration has been reviewed extensively and has important implications for prostate disease (44). Lineage tracing experiments demonstrate that multipotent basal epithelia give rise to basal, luminal, and neuroendocrine epithelia during prostate organogenesis (6, 16). In the homeostatic adult mouse, basal and luminal epithelia are largely maintained by unipotent progenitors (17, 36). Under castrate conditions, the prostate regresses due to the loss of luminal epithelia, but can be re-grown to its original size with androgen re-administration (18, 29). Select enrichment of castration-insensitive subpopulations characterized by expression of NKX3.1 (20), LGR5 (41), BMI1 (21), Sca-1 (22, 31), TROP2 (24), CD133 (23), LY6D (25), and KRT4 (26) have been described using lineage tracing, flow cytometry, and facultative regeneration assays. Label retention by slow cycling cells after multiple rounds of castration has also been proposed as a characteristic of stemness, and a transcriptional profile of labeled cells demonstrates survival of a luminal cell population after androgen deprivation (43). Castration-insensitive luminal epithelia that display many of these individual characteristics are shown to be enriched in the proximal region of the prostate, but defining a cell type as prostate based on ‘luminal’ (e.g. KRT8/18 expression) characteristics and anatomical location does not take into account the many different types of luminal cells that exist in this complex region where the prostate, ejaculatory ducts, and urethra merge. This is especially important to take into account when indelibly labeling KRT8+ cells with a fluorophore prior to prostate budding, where even the urogenital sinus epithelia marked by expression of KRT4 (Figure 4) also express KRT8 (6). Through scRNA-seq, we have been able to specifically identify a KRT8+ luminal cell type of the urethra that is the castration-insensitive progenitor of the proximal prostate described in previous studies (20, 22-26, 29-31, 39-43).

Single cell RNA sequencing provides an objective identification of cell types based on the expression of a wide range of transcripts. We previously used scRNA-seq to produce a cellular anatomy of the normal adult human prostate and prostatic urethra, revealing the existence of two novel urethral luminal epithelial cell types (club and hillock) that extend into the proximal ductal network that connects distal prostate glands to the prostatic urethra (4). In our own dissections of the mouse prostate lobes away from the urethra, we observe that 0.44 ± 0.10% of the total epithelia in the homeostatic proximal prostate are TROP2+ urethral luminal cells (Figure 3B), and that these cells are highly enriched in prostates from castrated mice (Figure 3C, D). Comparing the transcriptomic signature of Sca-1+ cells enriched in castration from a separate study (26) revealed high correlation with our urethral luminal epithelia (Figure 2D), suggesting that these are the same cell type. These data suggest that dissections of prostate lobes include extensions of the urethra into the proximal ductal network.

The incidence of benign and malignant prostate disease in proximal (transition/central zones) and distal (peripheral zone) anatomical regions, respectively, has long perplexed prostate biologists (9, 45). It has been proposed that BPH is a reawakening of morphogenic activity by prostate stem cells (13, 15), but the full identity of cell types that propagate prostate growth has been elusive. Given the dramatic increase in urethral epithelial cells in the prostates of 5ARI-treated men (Figure 5), it is tempting to postulate that BPH is a process of budding and branching of the urethra and that 5ARI treatment is ‘trimming the leaves’, but 3-dimensional reconstructions of the prostate ductal architecture will be necessary for confirmation. As possible evidence that BPH is a reawakening of urethral budding and branching, we demonstrate that the identity of urethral luminal epithelia is established prior to prostate budding in the developing mouse and human (Figure 4). Evidence from lineage tracing studies of mitochondrial mutations in humans demonstrates a clonal expansion of cells in the proximal urethra to the distal prostate in healthy adults (27, 28), but needs confirmation in BPH specimens.

Identification of urethral luminal cells as the castration-insensitive progenitor enriched in facultative stem cell assays could be interpreted in multiple ways. It is possible that the fetal mesenchyme used in facultative regeneration assays used to assess stemness is unnaturally inductive, as historical studies have shown that even adult bladder or urethral epithelium can be transdifferentiated to prostate (39, 46, 47). Moreover, spheroid and label-retaining assays might enrich multiple cell types that cycle relatively slower or survive better under certain *ex vivo* conditions than other cell types. Alternatively, urethral luminal epithelia (or even ejaculatory duct luminal epithelia, which express high levels of the putative stem cell marker CD133 (40)) are actually progenitors for the prostate under stressed conditions such as castration, inflammation, and cancer. These possibilities need to be assessed with tissue- and cell type-specific lineage tracing models.

Finally, it will be important to characterize the functional role of urethral luminal epithelial cells in maintaining the health of the prostate. Club and hillock epithelia of the lung are anti-inflammatory, anti-bacterial, and anti-viral and can also serve as progenitors in certain situations (8, 48, 49). The transcriptomic signature of urethral epithelia in the human and mouse prostate also suggests progenitor and immunomodulatory activity, suggesting the potential for a similar functional role in protecting distal prostate tissue from environmental exposures and infections (Figure 5C). The increased abundance of urethral epithelia within glandular nodules in BPH patients could simply indicate an expansion in response to infection or inflammation. Alternatively, these cells could be acting as progenitors that differentiate into prostate and contribute to growth due either to intrinsic properties of stemness or extrinsic signaling from the surrounding stroma (42).

## Acknowledgments

We thank the families of organ donors at the Southwest Transplant Alliance for their commitment to basic science research. We acknowledge the assistance of the University of Texas Southwestern Tissue Resource, a shared resource at the Simmons Comprehensive Cancer Center, which is supported in part by the National Cancer Institute under award number 5P30CA142543. We acknowledge the assistance of UT Southwestern’s Children’s Research Institute Flow Cytometry Core and the Whole Brain Microscopy Core. Financial support came from TL1TR002375 and F30DK122686 (H.M.R.), R01 DK115477 (D.W.S.); U54DK104310 (D.W.S. and C.M.V.), U01DK110807 and R01DK099328 (C.M.V); and the generous donations of the Smith, Penson, and Harris families to the UTSW Department of Urology (C.G.R.).

## Methods

### Contact for reagent and resource sharing

Further information and requests for resources and reagents should be directed to and will be fulfilled by the Lead Contact, Douglas Strand, following MTA approval.

### Human subjects

Healthy prostate specimens used in this study were obtained from 15, 17-42 year old male organ donors whose families were consented at the Southwest Transplant Alliance from March 2017 to January 2020 under IRB STU 112014-033. BPH specimens were obtained fresh from 28 patients undergoing simple prostatectomy at UT Southwestern Medical Center. Human lower urinary tracts were from the University of Pittsburgh Medical Center and Joint MRC / Wellcome (MR/R006237/1) Human Developmental Biology Resource (www.hdbr.org) under an approved University of Wisconsin-Madison IRB protocol (2016-137 0449). Clinical details for each fetal, normal adult, and diseased adult human specimen and their usage in associated figures are shown in Dataset S3.

### Mouse tissue collection and castration

Male C57Bl/J mice were obtained from the UT Southwestern Mouse Breeding core. Lower urinary tract tissues were collected from mice after euthanasia. Eleven week old mice were subjected to castration surgery following approved surgery protocols. The prostate was allowed to involute for 2 weeks before the mice were euthanized for tissue collection.

### Tissue processing

Fresh tissue samples were transported in ice-cold saline and immediately dissected into portions for fixation in 10% formalin followed by paraffin embedding. For human specimens, a 4 hour enzymatic digestion into single cells was performed at 37°C in 35 mls HBSS containing 5 mg/ml collagenase type I (Life Technologies), 10 μM ROCK inhibitor Y-27632 (StemRD), 1 nM DHT (Sigma), 1 mg DNAse I, and 1% antibiotic/antimycotic solution (100X, Corning) (4). Mouse specimens were digested for 1 hr in HBSS containing either 10 mg/ml cold protease or 1.5-2 mg/ml collagenase type I, plus 10 μM ROCK inhibitor Y-27632, 1 nM DHT, 1 mg DNAse I, and 1% antibiotic/antimycotic solution.

### Flow cytometry

Human and mouse prostate and prostatic urethral cells were analyzed in the UT Southwestern CRI Flow Cytometry Core on a BD FACSAria FUSION SORP or a BD FACS Aria II SORP 5-laser flow cytometer and analyzed with FlowJo software as previously published (4). Improved antibody panels based on single cell data were built with fluorescence minus one experiments. Dataset S4 displays information on antibodies used for flow cytometry.

### Immunohistochemistry

In brief, 5 μm paraffin sections were deparaffinized in xylene and hydrated through a series of ethanol washes. To block endogenous peroxidases, tissues were blocked with 0.3% H_2_O_2_ in Methanol for 20 mins. Following a wash in PBS, heat mediated antigen retrieval was performed by boiling slides in Vector antigen unmasking solution (Vector labs, H-3300) for 20 min in a conventional microwave oven. Tissues were blocked with 2.5% horse serum for 20 mins. For the first stage of staining, the first primary antibody diluted in 2.5% horse serum was applied for 1 hr at room temperature. Following washes in PBS, tissues were incubated with enzyme conjugated secondary antibody solution for 30 mins at room temperature. Tissues were washed twice in PBS and substrate solution was added to develop antibody stain. This was repeated for the second primary antibody. Horseradish peroxidase and alkaline phosphatase enzyme systems from Vector laboratories were used to obtain dual stains. Tissues were counterstained with hematoxylin and mounted with Permount solution. For primary and secondary antibody information see Dataset S4.

### Immunofluorescence

Fluorescent immunohistochemistry was performed as described previously (Henry et al. 2018). In brief, 5 μm paraffin sections were deparaffinized in xylene and hydrated through a series of ethanol washes. Heat-mediated antigen retrieval was performed by boiling slides in Vector antigen unmasking solution (Vector labs, H-3300) for 20 min in a conventional microwave oven. Tissues were washed with PBS and non-specific binding sites were blocked for 1 hr in blocking buffer (1x TBS, 5% Normal Horse Serum, 0.1% Bovine Serum Albumin, 0.1% Tween 20, 0.2 mM Sodium azide). Tissues were incubated overnight at 4°C with primary antibodies diluted in blocking buffer. Tissues were washed several times in PBS and incubated with secondary antibodies diluted in blocking buffer for 1 hour at room temperature. Following several washes with PBS, tissues sections were incubated with 4’,6-diamidino-2-phenylindole, dilactate (DAPI) to visualize cell nuclei and mounted in phosphate buffered saline containing 90% glycerol and 0.2% n-propyl gallate. Images were obtained using a Nikon Eclipse Ti-U with NIS Elements software or a Zeiss Axioscan Z1 microscope. For primary and secondary antibody information see Dataset S4.

### Single cell sequencing

Four mouse prostates and three mouse prostatic urethra were used. In addition, three young human prostate specimens were used previously (4) were sequenced deeper (see Dataset S2). Single cell suspensions were loaded into the 10x Genomics Chromium Controller using the Chromium Single Cell 3’ Library and Gel Bead Kit v3 according to the manufacturer’s protocol. Briefly, 17,400 total cells of each sample were loaded on individual lanes of a Single Cell A Chip with appropriate reagents and run in the Chromium Controller to generate single cell gel bead-in-emulsions (GEMs) for sample and cell barcoding. Libraries were generated using 10x Genomics’ protocol. Libraries were pooled and submitted for sequencing on an Illumina NextSeq 500 in high output mode. 150 cycle flow-cells were used to sequence 26 cycles for read 1, 58 cycles read 2, and 8 cycles for the i7 index.

### Single cell sequencing data analysis

Data analysis was based on previously published analysis (4, 50), modified in the following ways:

#### Cellranger

Cellranger mkfastq and count (version 3.1.0) was used to demultiplex, combine sequencing runs on the same sample, and to call cells. The reference genomes used were GRCh38 and mm10 (10x Genomics version 3.0.0). These 10x Genomics tools were automated and parallelized using the UT Southwestern Bioinformatics Core Facility (BICF) pipelines (51, 52).

#### Cell filtering

Low quality cells were filtered out by consecutively filtering each sample individually based on UMI counts, percentage mitochondrial content (%mito), and number of genes (in that order). Filter thresholds were chosen dynamically for samples based on the distribution of the parameter. The upper bound of the UMI filter was determined by removing cells with UMI counts lower than the highest frequency bin (10 bins) and scaling the remaining cells’ UMI between 0 and 360 and applying the RenyiEntropy thresholding technique. The lower bound of the UMI filter was set to 200. The bounds to the %mito and gene number filters were determined similarly. High %mito threshold was determined using the Triangle filter on the rescaled (0-360) parameter, from cells with a greater than or equal value to the highest frequency binned %mito (100 bins). The lower bound to the gene count filer was determined using the MinErrorI filter on the rescaled (0-360) parameter, from cells with less than the value to the highest frequency binned gene count (100 bins). Cells were not filtered for low %mito or high %gene count. RenyiEntropy, Triangular, and MinErrorI thresholding was applied using autothresholdr version 1.3.5 (53).

#### Aggregation

Samples were aggregated by normalizing with the sctransform (version 0.2.0) method (54), and using Seurat’s (55, 56) (version 3.1.0) reciprocal PCA method. Samples with less than 750 cells (post-filter) were merged prior to aggregation.

#### Stressed cell removal

Cells displaying high stress signatures were removed. Aggregated cells were scored for stress using Seurat’s AddModuleScore method, using genesets enriched in stressed cells. For human samples, the geneset came from Henry, *et al.* (4), while the mouse geneset came from van den Brink, *et al.* (57).

#### Clustering

Principal component analysis (PCA) was performed on the data. Graph-based clustering was performed using the principal components which represents 90% of the cumulative variance associated (using Seurat).

#### Human cell type identification

The tool SingleR (58) (version 1.0.0) was used to identify cell types. Broad cell lineages were identified utilized the main labels from the built-in Human Cell Atlas data (59). This was done in cluster mode using a resolution of 0.5. Cell types were merged based on their membership in the following major groups: epithelia, fibromuscular stroma, endothelia, and leukocyte. The prostatic epithelial and fibromuscular stromal cell types used cell identities from Henry, *et al.* (4) in single-cell mode.

#### Human to mouse cell type transfer

Mouse cell types were identified similarly as the human. Broad cell lineages were identified using the main labels from the built-in mouse Immunological Genome Project data (60). The cell types were merged as with the human data. Epithelial cells were subset, and re-clustered (as above). Prostate epithelial cell types were labeled using the identified data from the human samples, with the genes converted to mouse orthologs (genes with no orthology were removed) in cluster mode, at a resolution of 0.1. The ortholog map was retrieved from Ensembl’s (61) BioMart from release 99 GRCm38.p6 data.

#### DEG calculation

Differentially expressed genes were calculated from the sctransform normalized data, using the MAST (62) method, implemented in Seurat. Significance determined using the Bonferroni corrected p-value, with an alpha of 0.05.

#### Clustering analysis

Three patient specimens dissected into transition/central zone and peripheral zone. Each zone was sorted for viability before loading into the 10x Genomics chromium controller. The 10x Genomics’ analysis pipeline, cellranger (version 2.1.1) was first used to demultiplex and produce a gene-cell matrix. Bcl files were demultiplexed using their barcode-aware wrapper for bcl2fastq (version 2.17.1.14). Transcriptomes were aligned to GRCh38 using STAR (version 2.5.1b). Samples were then aggregated by downsampling to match their mean mapped reads per cell. Low quality cell barcodes were filtered out using 10x Genomics’ algorithm (high quality barcodes = total UMI count ≥ 10% of the 99^th^ percentile of the expected recovered cells). Dataset S2 displays the sequencing metrics for each barcoded experiment.

Seurat (version 2.3.1), an R toolkit for single cell transcriptomics formed the basis of further analysis (55) run on R version 3.4.1. Genes that were expressed in three cells or less were filtered out along with cells expressing fewer than 200 unique genes. Cell cycle state was predicted based on Seurat’s built in principal-component (PC) analysis. Briefly, cells were scored based on expression their expression of G2M and S phase genes (63). Low quality cells and multiplets were excluded by removing cells with fewer than 500 unique genes and greater than 3,000 unique genes, as well as cells with greater than 10% of their transcriptome being mitochondrial genes. Data was then scaled to 10,000 and log transformed. Mitochondrial genes were then removed from further analysis. UMI counts were then scaled and variation due to differences in UMI/cell, percent mitochondrial genes, and cell cycle phase were regressed out of the data using a built-in Seurat function. Cells from the three patients were then subsetted and recombined using canonical correlation analysis (CCA) in order to align the clusters. The highest variable genes were found with an algorithm developed by Macosko *et al* (64), and were defined as an average expression between 0.2 and 5 with a dispersion greater than 1. The intersection of these genes between the three patients were used to calculate 50 CCAs and the first 30 were aligned. These 30 aligned-CCAs were used for t-SNE visualization and clustering. Cells were clustered using a graph-based clustering approach (55). Briefly, cells were embedded in a KNN graph structure based on their Euclidean distance in PC space, with edge weights refined by shared overlap in their Jaccard distance. Different resolutions were generated based on a granularity input.

#### DEG calculation

Cluster/cell identities were aggregated and DEGs for the identified cell types were then determined using a Wilcoxon rank sum test on genes present in at least 25% of either the population of interest or all other cells in that lineage which are at least two-fold enriched in the population of interest and a maximum Bonferroni corrected p-value alpha of 0.05.

#### Geneset Enrichment Analysis

Differentially expressed genes were calculated E-MATB-4991 (26) microarray data using Transcriptome Analysis Console using default setting (SST-RMA). Each sample type’s DEGs were calculated compared to the other two groups (alpha = 0.05). These DEGs were used to correlate to the expression of the identified mouse epithelial cell types using the tool QuSAGE (version 2.20.0).

#### KEGG Analysis

Enrichment of KEGG annotated genesets was calculated using DAVID (version 6.8) from DEGs up in human BPH club cells compared to human normal club cells. Significant genesets were determined using an alpha of 0.05 on Benjamini corrected p-values.

### Data and software availability

To increase rigor, reproducibility and transparency, raw image files (including duplicates not displayed here) and scRNA-seq data generated as part of this study were deposited into the GUDMAP consortium database and are fully accessible at: https://doi.org/10.25548/16-WM8C (65). The scRNA-seq data from human tissues were deposited into the GEO series GSE145843 (normal) and GSE145838 (BPH), while mouse scRNA-seq data can be found in GEO series GSE145861 (prostate) and GSE145865 (urethra). R code used to produce all the scRNA-Seq analysis can be found at https://git.biohpc.swmed.edu/StrandLab/sc-TissueMapper.git (tag: 2.0.0) (66). Analyzed scRNA-seq data from mouse and human specimens can be found at http://strandlab.net/sc.data, where gene expression can be investigated in the cell type clusters identified in this study.

## Supplementary Information

**Figure S1.**
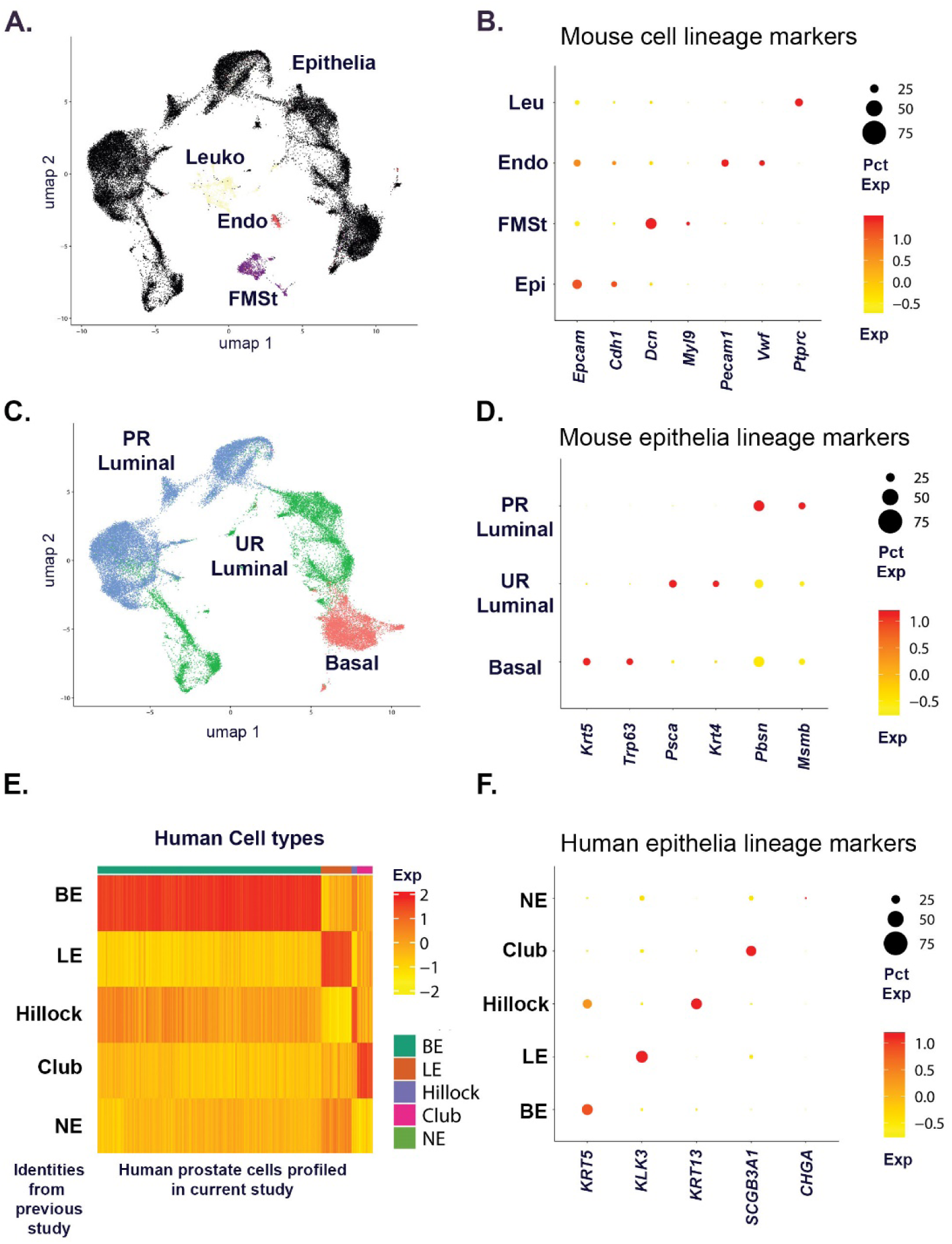
Identification of mouse prostate and urethral cell types. **(A)** Aggregated scRNA-seq data from mouse prostate and urethra (n=3) with major lineages identified using human prostate data (n=3). **(B)** Anchor genes for each lineage displayed by dot plot. **(C)** Sub-clustered epithelia with clusters identified using human prostate scRNA-seq data. **(D)** Anchor genes for each epithelial lineage displayed by dot plot. **(E)** Correlation of human prostate epithelial cell signatures from previous study with current data set used to establish mouse epithelial cell identities. **(F)** Dot plot of cell type-specific genes used from human prostate cell type signatures.

**Figure S2.**
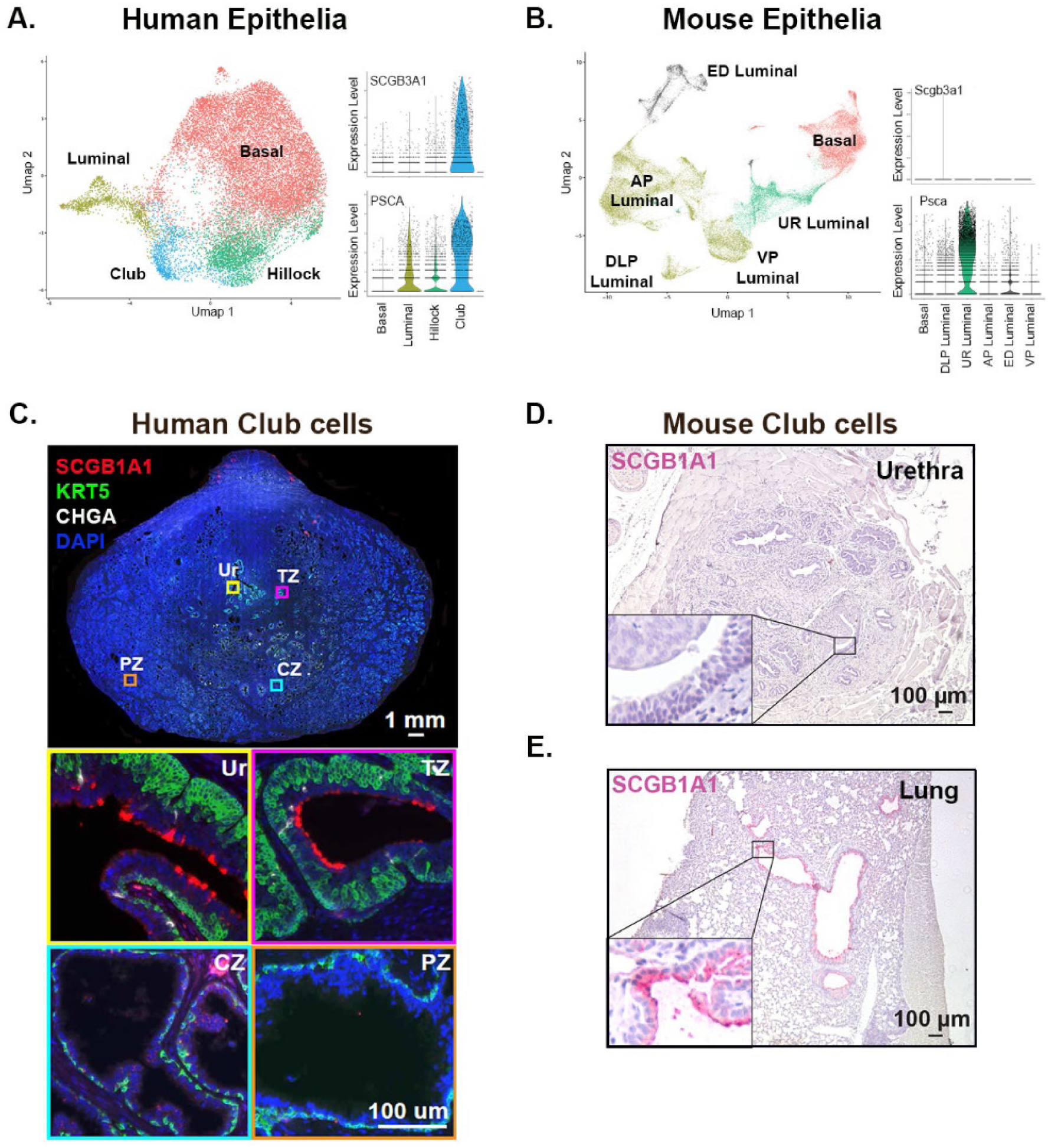
Differences in urethral luminal epithelia in mouse vs. human. **(A)** UMAP plot of previously published human prostate and prostatic urethra scRNA-seq with adjacent violin plots of club cell (SCGB3A1) and pan-urethral luminal cell (PSCA) markers. **(B)** UMAP plot of mouse prostate and urethra scRNA-seq with adjacent violin plots of club cell (SCGB3A1) and pan-urethral luminal cell (PSCA) markers. **(C)** Immunofluorescence of young adult prostate highlighting localization of SCGB1A1+ club cells in the prostatic urethra and proximal ducts. **(D)** Immunohistochemistry of mouse urethra with club cell marker SCGB1A1 showed no reactivity while reactivity was seen in the mouse lung as a positive control **(E)**.

**Figure S3.**
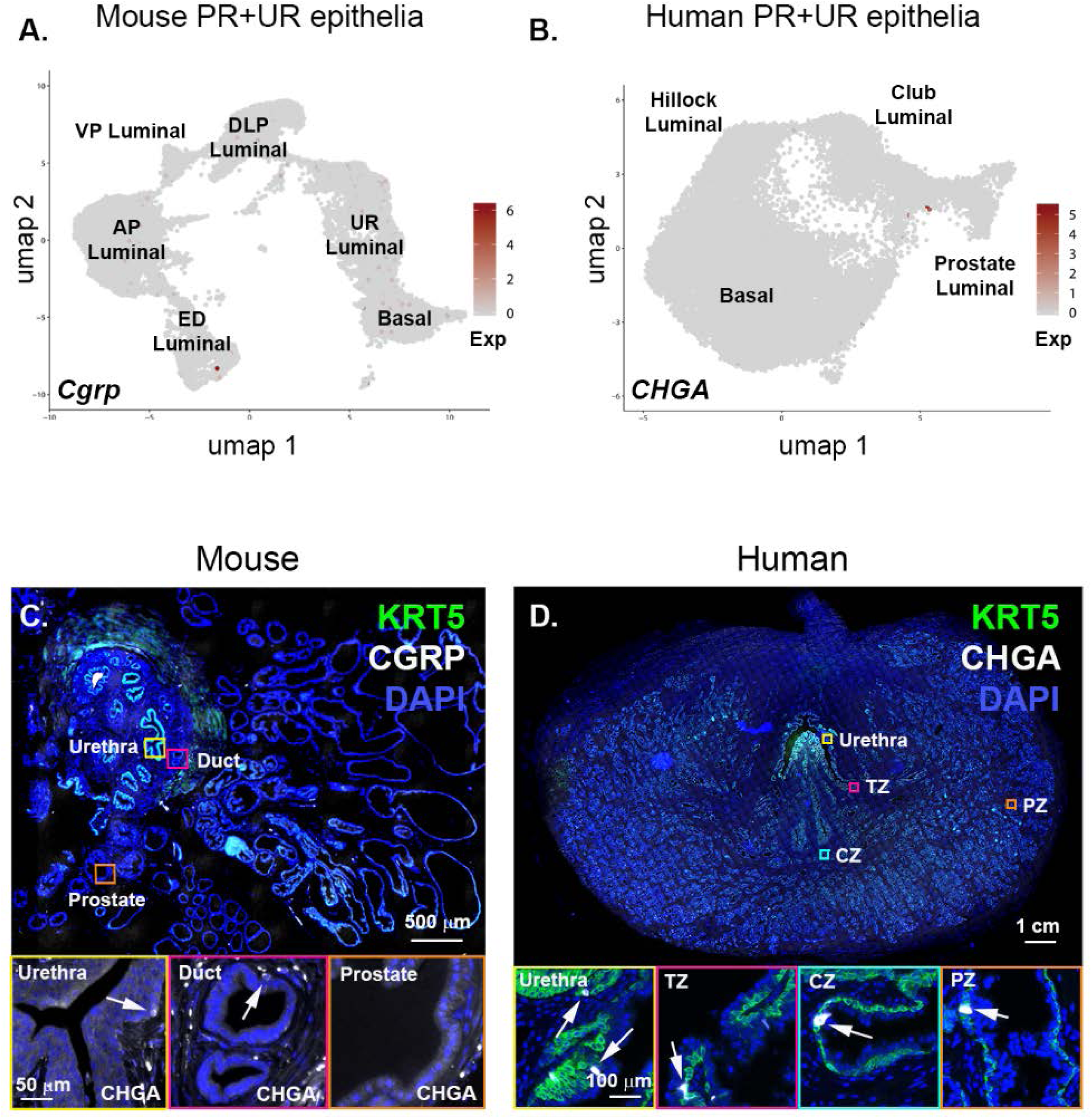
Cognate neuroendocrine epithelia (NE) of the mouse and human prostate. **(A)** UMAP plot of mouse prostate and urethral epithelia with NE highlighted with CGRP expression. **(B)** UMAP plot of human prostate and urethral epithelia with NE highlighted by CHGA expression. **(C)** Whole mount transverse mouse lower urinary tract with basal (KRT5) and NE (CHGA) cells highlighted by immunofluorescence. KRT5 layer is removed in insets to better visualize NE cells. **(D)** Whole mount transverse human normal prostate with basal (KRT5) and NE (CHGA) cells highlighted by immunofluorescence. PR, prostate; UR, urethra.

**Figure S4.**
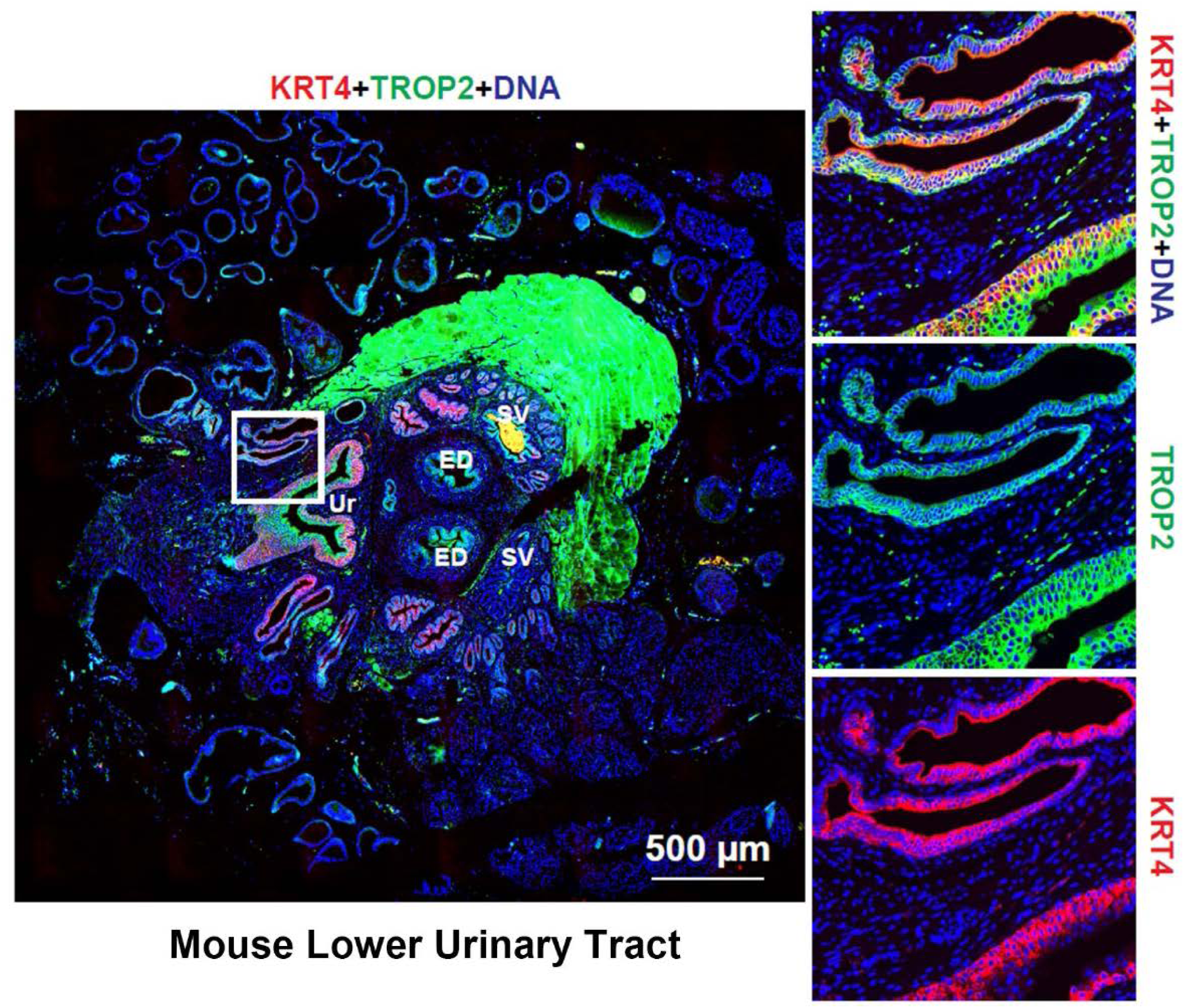
TROP2+ urethral luminal epithelia are co-express KRT4.

**Figure S5.**
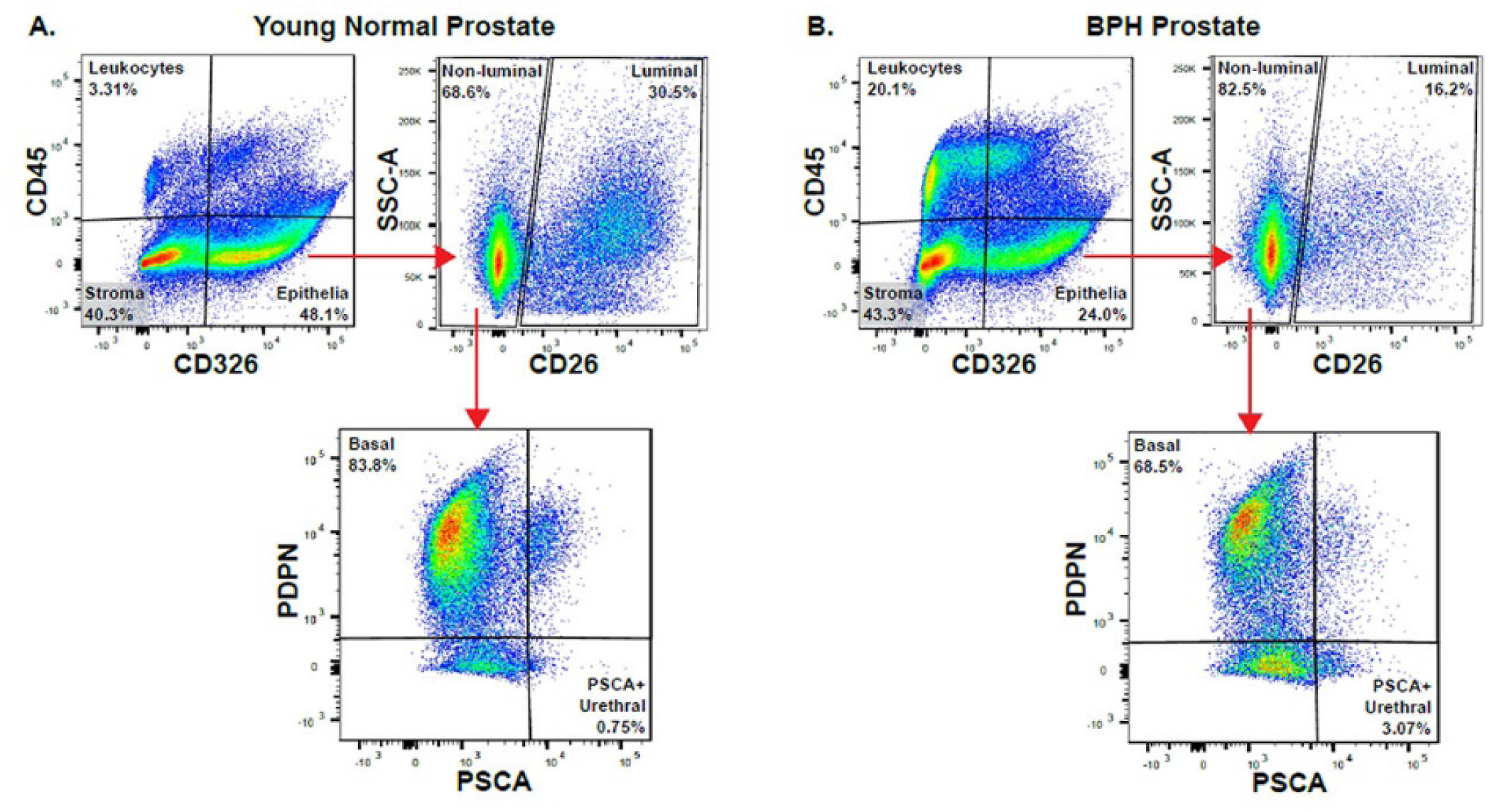
Flow cytometry scheme for quantitation of urethral luminal epithelia in normal vs. BPH human prostate. Panels representative of n=6 independent samples/group.

**Dataset S1 (separate file).** Differentially expressed genes for each mouse cluster.

**Dataset S2 (separate file).** Single cell RNA sequencing metrics for each specimen.

**Dataset S3 (separate file).** Clinical data for each human specimen.

**Dataset S4 (separate file).** Antibody information for flow cytometry and immunostaining.

**Dataset S5 (separate file).** Differentially expressed genes for each human prostate and urethral epithelial cell type in normal vs. BPH.

## Notes

https://www.strandlab.net

